# Bridging the Python Training Gap for Bioscientists in Brazil: Improvements and Challenges

**DOI:** 10.1101/2024.11.25.624749

**Authors:** Gustavo Schiavone Crestana, Ubiratan da Silva Batista, Michelli Inácio Gonçalves Funnicelli, Larissa Graciano Braga, Luíza Zuvanov, Rodolfo Bizarria, Raissa Melo de Sousa, Pedro Henrique Narciso Ferreira, Flavia Vischi Winck, Gabriel Rodrigues Alves Margarido, Alessandro de Mello Varani, Diego Mauricio Riaño-Pachón, Renato Augusto Corrêa dos Santos

## Abstract

The rapid evolution of high-throughput technologies in biosciences has generated diverse and voluminous datasets, requiring bioscientists to develop data manipulation and analysis skills. Python, known for its versatility and powerful libraries, has become a crucial tool for managing these datasets. However, there is a significant lack of programming training for bioscientists in many countries. To address this knowledge gap among scientists in Brazil, the Brazilian Python Workshop for Biological Data was introduced several years ago, focusing on basic programming concepts and data handling techniques using popular Python libraries. Despite the progress and positive feedback from earlier editions, challenges persisted, necessitating continuous adaptation and improvement to meet the evolving needs of bioscientists.This work describes the advancements made in the 2021 and 2022 editions of the workshop and discusses new suggestions for its ongoing enhancement. Key innovations were introduced in the workshop structure and coordination, including the creation of new committees and the establishment of a code of conduct. Feedback forms were updated to enable real-time adjustments during the event, improving its overall effectiveness. The workshop also expanded its reach by increasing geographical diversity among participants. New didactic strategies, such as pair-teaching, code clubs, and the integration of information and communication technologies (ICTs), were implemented to enhance learning outcomes. Programming best practices and scientific reproducibility were emphasized through talks and hands-on activities, guided by PEP8 conventions. Furthermore, efforts to enhance scientific dissemination were intensified, with an increased presence on social media and participation in international scientific events and communication networks. Finally, we present updated recommendations for students, researchers, and educators interested in organizing and promoting similar events, building on those previously described.

## Introduction

In recent years, bioscience has undergone a data revolution due to advances in high-throughput technologies such as genomics, transcriptomics, proteomics, and phenomics driving progress not only in medicine, agriculture, and environmental science, but also in numerous related fields. The availability of massive amounts of biological data has created unprecedented opportunities for scientific discovery [1,2]. Alongside that, other types of data, such as georeferencing, biometric, and phenotypic, are increasingly being generated and analyzed in bioscience research [3].

Overall, the increasing availability of different types of biological data has opened up new avenues of research. The analysis and interpretation of large and complex datasets is often challenging, and bioscientists are required to develop skills in data manipulation, statistics, bioinformatics, and programming [4,5]. This “new kind of scientist” [6], capable of bridging life sciences and computer science knowledge is rare, demonstrating an important bottleneck in modern bioscience [7,8].

The Python programming language has become popular as a tool for manipulating biological data due to its versatility, user-friendly feel, and availability of powerful libraries aimed at automating analyses and data visualization [9]. In addition to the ability to manipulate large datasets, the reproducible nature of the open-source language makes it an attractive tool for areas of life sciences, which continue to generate and analyze large volumes of data [10]. Despite the growing importance of computational programming in handling biological data, the lack of training initiatives for bioscientists is notable, such as in Brazil and other Latin American countries. Despite the country’s significant contributions to bioscience research, Brazilian scientists generally still lack the necessary skills to effectively analyze and interpret data sets generated by modern bioscience research [11,12]. Although in recent years, some Brazilian introductory training initiatives focused on manipulating biological data using Python have emerged in different areas of knowledge (S1 Table), it is surprising how limited these initiatives still are, given the potential of the Python language to enhance bioscience research and improve scientific outcomes.

In response to this lack of training, we developed an initiative to teach introductory programming skills to bioscientists. The Brazilian Python Workshop for Biological Data was first held in 2017 and was previously described by our group [13]. The workshop aimed to provide participants with the necessary skills to handle and analyze large biological datasets using Python. In the 2020 edition, the course consisted of two modules: the first covered basic programming concepts such as variables, data types, loops, and conditional statements, while the second focused on data manipulation and visualization using Python libraries such as Biopython [14], Matplotlib [15], Numpy [16], and Pandas [17]. The course was delivered using a mix of live coding, lectures, and hands-on exercises, and it was focused on bioscientists with little to no prior programming experience [13].

Inspired by the work developed by The Carpentries initiative [18], gathering feedback from our community was necessary to understand and accommodate their needs and motivations [19]. In this sense, our previous work had positive results, as participants reported significant improvements in their programming skills and a greater ability to analyze biological data using Python [13]. Furthermore, with the experiences gained during the initiative, we previously recommended the use of open-source software and free online resources to make the course more accessible to a wider range of participants, particularly those from low-income backgrounds. We also emphasized the need for a diverse group of instructors, with a range of expertise in both programming and bioscience, to ensure the course material was relevant and applicable to different research fields. Feedback and perspectives from previous editions provided suggestions to further improve the initiative based on our experiences (in Box 2 from [13]).

Overall, the recommendations in our previous study highlight the importance of continuous improvement and adaptation of training initiatives to meet the evolving needs of bioscientists in data manipulation and analysis using Python. Also, some considerations focused on the organization and institutional aspects of the event. Finally, they endorsed the creation of a practice community for participants to continue learning and collaborating after the course had ended.

New editions of the Brazilian Python Workshop for Biological Data were held virtually in 2021, 2022, and 2023, trying to incorporate the suggestions and recommendations made by [13]. In this context, this manuscript describes the advances made during two of the most recent editions, 2021 and 2022, and discusses new suggestions for the continuous improvement of a workshop focused on bioscientists with little or no programming experience.

## Results

### General characterization of the 2021 and 2022 editions

To accurately portray and characterize the participants in the workshop editions held in 2021 and 2022, feedback forms were applied throughout the event, containing inquiries pertaining to geographical, ethnic, gender, academic, and professional diversity (https://github.com/brazilpythonws/wpbd_update_2021_2022). Although a new edition with a similar format occurred in 2023, the data from this latest edition was not included in this manuscript. To understand the target audience and key aspects of the course, as well as to pinpoint challenges, highlights, and recommendations for future events, we collected and analyzed data using both objective and open-ended questions. The organizing committee utilized fundamental Python programming concepts to create informative visual representations based on the question type, data acquired, and specific Python libraries employed (Table 1).

**Table 1.** Graphic output generated by the analysis of collected data from feedback forms of both editions.

| Question type | Data type |  | Used library | Graphic output |
| --- | --- | --- | --- | --- |
| Objective | Qualitative | Nominal | Basic Python Functions/Matplotlib | Donut/Bar chart |
|  |  | Ordinal |  |  |
| Open-ended | Qualitative | Ordinal |  | Stacked Bar Chart |

### Characterization of registrations

Comparatively, an increase in enrollment was reported from the first edition (2017) until 2022 edition (Fig 1A). A peak was observed during the first virtual edition in 2020 with 350 registrations. Because 2020 was the year of the COVID-19 outbreak and the need for social distancing and restrictions on large gatherings, many organizations and event planners opted for virtual events as a way to continue their activities while keeping participants safe [20,21]. The pandemic has accelerated the trend of virtual events, and many organizations are likely to continue using this format in the future. As a result, there was a notable increase in the number of virtual events [20], possibly explaining the higher enrollment rate seen in Fig 1A. After 2020, there was a decrease in registrations (2021), but the number rose again for the 2022 edition with more than 300 registrations.

**Fig 1.**
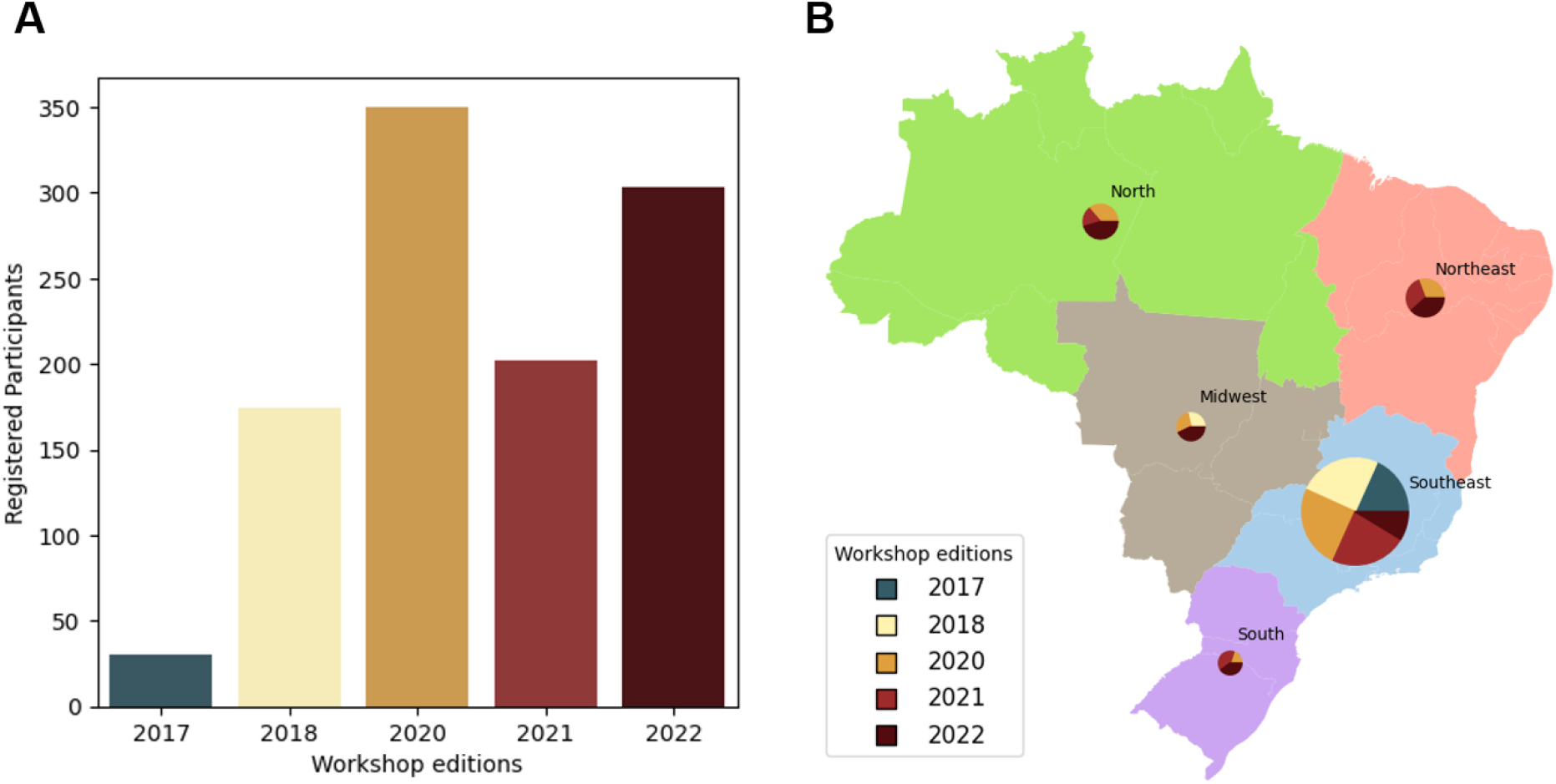
Evolution of number of participants and registrations in the Brazilian Python Workshop for Biological Data. (A) The number of registrations across the editions. The peak in registrations during the 2020 edition (the first conducted in a virtual format) can likely be attributed to the implementation of social distancing measures and restrictions on large gatherings due to the COVID-19 outbreak; (B) The pie charts overlaid on the map of Brazil illustrating the expanding geographic distribution of participants selected in the workshop over the years; **Note:** The cumulative total of participants selected over the years was 140, distributed as follows: Southeast (104), Northeast (13), North (11), Midwest (7), and South (5).

For these editions, 37 (2021) and 40 (2022) participants were selected according to predefined criteria, based on previous recommendations [13]: (I) little or no prior knowledge of the Python programming language, (II) analysis of students’ research activities and their academic involvement and (III) analysis of students’ expectations about the course. To create an egalitarian environment, the gender ratio among selected participants was kept proportionately equivalent (approximately 50% each).

An important aspect of the workshop conducted virtually was the deliberate inclusion of participants from all geographical regions of Brazil. In the 2021 and 2022 editions, an intentional effort to increase the number of participants from the Northeast, North, and Midwest regions was perceived in order to ensure a more equitable representation of all geographic regions across the country (Fig 1B).

### Workshop structure

Similar to the 2020 edition, both subsequent editions were structured online with the introduction of data collection, organization, and analysis using Python through practical sessions, employing a participatory live coding format [22]. We designed both workshops to take place in five days of immersive Python learning, with a one-day break for exercises (S2 Table and S3 Table). They encompassed introductory sessions covering language syntax concepts and practical classes involving Python coding and programming logic. Lectures were developed using interactive Jupyter notebooks hosted in the Google Collaboratory platform (https://colab.research.google.com/). During problem-solving and presentations, instructors actively wrote and narrated the code for the students. We encouraged the students to write, document, and run their own code independently and concurrently. The full description of both editions can be found in the repository link: https://github.com/brazilpythonws/wpbd_update_2021_2022.

### Keeping the workshop up to date: Following recommendations from previous research and creating innovative approaches

We provided a set of recommendations for those planning to organize similar workshops ([13] - Box 2), including activities implemented in the 2020 online edition and also directions for future workshops (i.e. regarding the format, virtual or in-person). In this section, we summarized advancements achieved during two recent editions of the workshop and in the next section, we discussed these advancements based on a temporal perspective.

#### 1. Cracking the code: Unveiling the *a priori* organizational structure of the workshop

The workshop success was due to a dedicated and diverse organizing team, whose roles extended beyond instruction to include social media management, ethical committees, and course material support. Despite minor changes, the core team structure remained consistent from 2021 to 2022 (Fig 2). Key innovations included a Coordination team, composed of at least one member from all workshop committees, serving as a deliberative body and providing overall supervision for the event. Secondly, the Communication and Media and the Scientific committees were established for event dissemination and overseeing participant management, respectively. Thirdly, a code of conduct (https://github.com/brazilpythonws/wpbd_update_2021_2022) was established to outline the principles adopted in the workshop for both organizers and participants to ensure mutual respect and prevent conflicts [23]. The event discouraged intolerance towards cultural, religious, funding conditions, sexual, and gender orientations, promoting diversity. Finally, a simplified Train-the-Trainers-like meeting [24], a brief virtual gathering involving former organizers (or advisers) and current committee members, was implemented to provide training and guide the new committee members. Feedback forms were developed to learn more about the participants and assess their contributions to the Workshop. Daily feedback was used as a basis for small-scale adjustments during the event, such as a quick review of a specific topic. The Organizing Committee carefully considered students’ complaints and modifications proposed in the final event document form (S4 Table and S5 Table).

**Fig 2.**
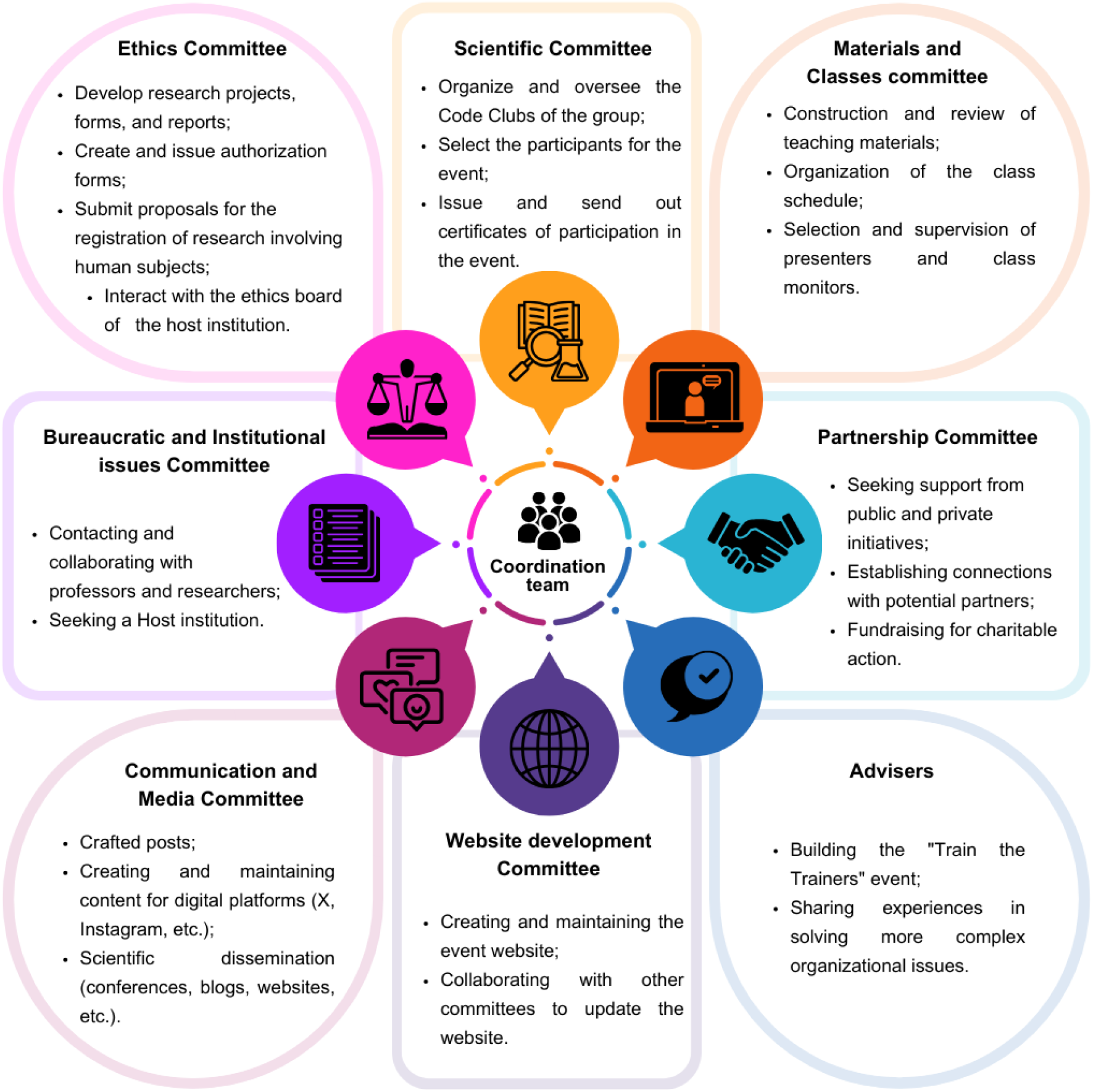
Workshop core structure in recent editions. Enhancements to the workshop’s core structure in two of the recent editions include detailed workload allocation among various committees, alongside adviser support, for organizing both the 2021 and 2022 editions of the Brazilian Python Workshop for Biological Data.

#### 2. Expanding access: democratization within the workshop

Due to the social distancing measures imposed by COVID-19, since 2020 the workshop shifted to an online format, allowing broader participation from various locations in Brazil (Figure 1).

Furthermore, in order to make the event more diverse and democratic, participant selection was carried out blindly since 2021. Recruiters had access only to the candidate’s answers, thereby reducing potential discrimination. Also, the proportion of men and women was maintained among the participants.

Additionally, data regarding other social aspects beyond gender, such as ethnicity, and any form of disability, were collected (although these aspects were not considered for the selection process) through virtual forms. Accessibility for students with disabilities was emphasized, with questions on disabilities and support needs included in those forms (https://github.com/brazilpythonws/wpbd_update_2021_2022). Although the event format and implemented methods are not yet sufficient to ensure total accessibility, they have allowed significant progress in building a more diverse community. Graphic output generated by the analysis of collected data from feedback forms of both editions also featured inclusive color palettes (https://github.com/brazilpythonws/wpbd_update_2021_2022), with plans for further accessibility enhancements like text-to-speech and sign language in future editions.

#### 3. Didactics: Recent editions highlights

Python and R are favored for teaching biomedical scientists due to their efficiency in skill acquisition [4]. Our workshop focuses on teaching Python using active methods like problem-solving [25], case-based approaches, live coding, individual and group exercises, and instructor interaction. In the 2022 edition, we introduced a teaching approach (hereafter called pair-teaching) where we combined an experienced and a beginner instructor as part of a strategy focused on sharing expertise among them, and training new instructors. Also, we established ‘Code Clubs’ for in-depth study (S1 Text), where the organizing team met twice a month to deeply study Python programming language and its primary applications for biological data. The workshop also considered individual, social, and cultural aspects for optimal learning [26], using daily feedback and event evaluations for continuous improvement [27]. Challenges like technical issues and technology access were minimized and monitored. We used interactive platforms like Discord, Google Meet, and Google Colab, along with *Flash Talks* and group exercises, to enhance student involvement [28]. However, we faced challenges such as the implementation of new Information and communication technologies (ICTs), which are further discussed.

#### 4. Programming best practices and scientific reproducibility

In order to introduce concepts about programming best practices, we included specific talks and hands-on activities in both editions. Furthermore, we initiated discussions on concepts such as reproducibility and readability. These topics help prevent inadequate code design and errors, which can lead to serious consequences [29,30]. We encouraged using well-established libraries, reading documentation, avoiding copy-pasting, using good editors, and testing code. We also promoted good practices for maintenance, debugging, and code sharing, such as using PEP8 conventions (https://github.com/brazilpythonws/wpbd_update_2021_2022), adding comments and writing documentation [31]. The workshop supports reproducible and shareable science, with materials from 2021-2022 available online in English (https://github.com/brazilpythonws/wpbd_update_2021_2022). Additional materials were shared through social media (see next section).

#### 5. Helping others to grow: Scientific dissemination and social media

Through the creation of the media and communication committee, we expanded year-round interaction with the audience and the impact of the teaching and research activities developed. For this purpose, in 2021 and 2022 we intensified our efforts to disseminate content on our social media profiles: Instagram (https://www.instagram.com/brazilpythonws) and X (https://x.com/BrazilPythonWS?s=20). Posts focused on instructive content, innovation, and practical applications of Python in biological data were published weekly on Instagram. All materials resulted from discussions of academic articles by the organizing team, topics covered in the code clubs, and teaching materials related to the basic syntax of the language, among other subjects. The increased online activity favored strengthening the bond between the organization team and the community, consisting of participants, former participants, enthusiasts, and collaborators of the event over time. Since this committee’s creation, there has been a growing demand from the audience (both young and adult) for the content presented (S1 Fig), reflected in the increase in the followers on social media and the expanded reach of the shared content (S2 Fig).

As part of our scientific dissemination role, we presented our initiative at international scientific events [32] and scientific communication networks [33]. The increasing visibility has motivated us to encourage other initiatives with similar goals within our community. In this regard, the 2022 edition featured a fundraising campaign to support an organization that promotes scientific and Python programming education, with a specific focus on bioinformatics for women (https://pyladiesbioinfo.vercel.app/).

### Navigating challenges: Insights and recommendations for future workshops

In order to provide a temporal perspective of our workshop, Fig 3 encompasses three comprehensive topics that compile all activities carried out before, during, and after each edition of the event, subdivided by the edition year: 2017/2018, 2020, 2021/2022, and future workshops. Our primary goal was to emphasize innovations and novelties in comparison to previous years, showcasing what worked among the implemented initiatives and offering insights into future perspectives on the respective topic.

**Fig 3.**
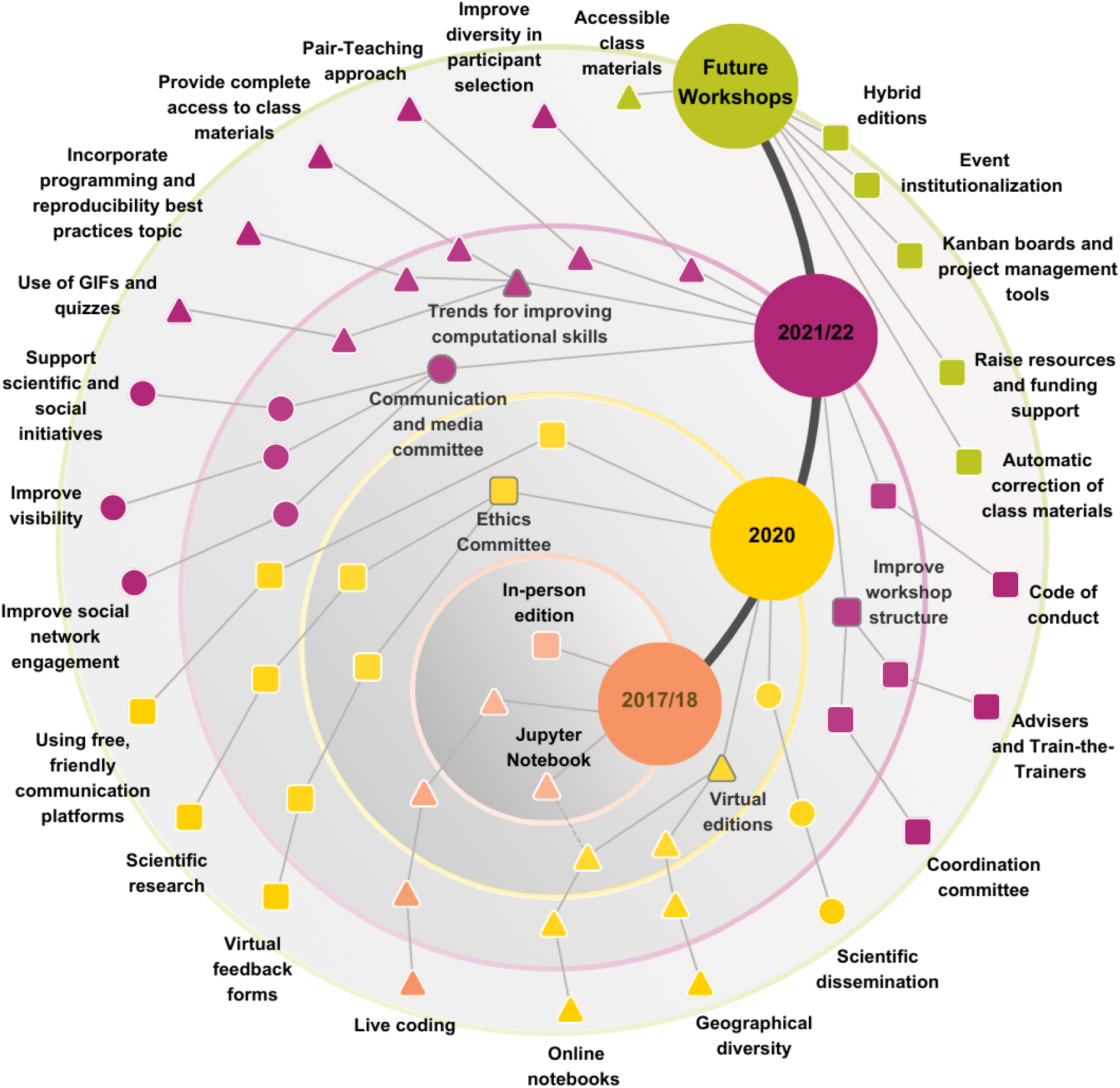
Evolution of temporal progress dynamics across successive workshop editions. Each node represents advancements implemented throughout all workshop editions or those under consideration for the future. Structural advancements (squares) denote impactful changes in the event’s structure and organization, encompassing both technical and bureaucratic aspects. Didactic advancements (triangles) highlight approaches fostering a more democratic environment conducive to knowledge transfer and consolidation. Lastly, communication and media enhancements (circles) denote improvements in the event’s social impact and interaction with its target audience and community. In this representation, each node’s color corresponds to the edition year of its inception, illustrating the evolution of advancements. Edges symbolize continuity across different editions, while unconnected nodes indicate advancements that were not carried forward into subsequent editions.

The first subsection, named “Collaborative forces in the workshop structure” (Fig 3 - structural advancements), elaborates on aspects concerning the coordination, technical components, institutional, and academic facets of the event. The second, labeled as “Enhancing educational strategies” (Fig 3 - didactic advancements), covers elements associated with the pedagogy and training of participants. Last, the third (“Engagement and social media Commitment”) highlights the progression of scientific outreach and the event’s social media dissemination (Fig 3 - communication and media enhancements). Here, we present and discuss all three of them:

#### 1. Collaborative forces in the workshop structure

Over time, the workshop framework has evolved into a more robust configuration, marked by an expansion in the number of organizers with previous experience in orchestrating similar events. Initially, participant groups were modest, but subsequent editions recruited new members from among the participants. The 2020 and 2021 editions were crucial for restructuring the framework, creating committees that reduced individual workloads. Tasks, once reliant on a few individuals, were delegated to thematic groups under coordination. The modified Train-the-Trainers-like program, advisers’ participation in meetings, and their inclusion in key discussions positively impacted the whole committee. Advisers, comprising ex-participants, offered suggestions and insights, saving time and preempting issues encountered in prior editions. The Train-the-Trainers-like initiative engaged former event members to conduct workshops, providing essential information on organization, logistics, common queries, and valuable tips. This structure, based on our experiences, could be functionally useful despite necessary modifications. Former students maintained the event’s continuity, crucial for building an engaged community, preserving the event’s identity, and providing valuable perspectives from personal experiences [34].

Transitioning to a virtual format allowed collaboration with a geographically diverse pool of individuals, enhancing accessibility for organizers and participants. Initially, communication relied on in-person meetings, but later editions used Slack and, in 2022, Discord due to changes in Slack’s free version policies. Future workshops should use collaborative platforms for preparation and interactive communication, considering project management tools like Trello and Kanban boards for better organization. Digital platforms managed by Google, such as Drive, Meet, and Colab, were selected for their integration, user-friendly interface, and compatibility. Since the 2020 edition, Google Colab was adopted since it does not require installation and runs entirely on the cloud. Before 2020, participants programmed using notebooks installed locally using Anaconda (an open source management system that installs, runs, and updates packages easily; https://docs.conda.io). Future editions will continue using these platforms, exploring tools like Google Classroom for creating and sharing grading assignments.

The workshop, hosted by various institutions and conducted online since the COVID-19 pandemic, ensured continuity and accessibility until 2023. This transition allowed participation from different regions of Brazil but introduced challenges like internet instability, virtual fatigue, and limited networking [13,35]. Returning to an in-person format introduces new challenges, such as adjusting to a new format and ensuring participation from different locations. Moving to a hybrid event could enhance interactions, engagement, and learning, offering networking opportunities and flexibility [20,36,37]. However, it also increases logistical complexity, requiring equipped facilities and careful planning [20]. Funding support is also an emerging need since substantial resources are required for logistics, infrastructure, and participation of both speakers and students. Despite the importance of having funding, our workshop has never charged a registration fee. In future workshops or initiatives, the development of effective fundraising strategies and the establishment of robust institutional partnerships are imperative for the mobility of the event to an in-person or hybrid format. Possible opportunities for funding support include outreach scholarships and public research agency funding calls.

Creating a proper learning environment involves not only structure but also expected conduct. Developing a code of conduct was prioritized, functioning as a behavioral guide to uphold good practices within the event’s community [38,39]. It took several editions for our event to formulate a robust Code of Conduct with inputs from the entire organizational team. However, discussions on this matter have consistently occurred and undoubtedly shaped the structure of our code throughout the editions.

Feedback from participants and organizers, collected through methods developed across editions, has been crucial for improvements. Initially inspired by events like Software Carpentry workshops (https://software-carpentry.org/), feedback was collected via post-it notes and later through Google forms, enhancing learning outcomes and skills practice by facilitating interactive role play and fostering collaborative decision-making [40]. Also, virtual feedback allows more frequent adjustment instruction based on data patterns by organizers [41]. Therefore, both the adoption of virtual forms and the establishment of the ethics Committee strongly influenced the format of feedback, data collection, and their analysis (S3 Fig). We recommend the continued use of virtual forms for reliable feedback, essential for rethinking strategies and implementing new ideas. Additionally, investigating new research avenues, such as the effectiveness of live coding for teaching programming, is crucial for advancing knowledge in our field.

Sharing workshop materials was improved by sharing digital notebooks and presentations on social media. Future editions aim to include materials in a permanent database with an object identifier (DOI) for accessible and open education.

During the workshop’s 2021 edition, we attempted to automate assigning and grading students’ notebooks using the nbgrader system, an open-source tool popular in educational contexts [42,43]. However, auto-grading tools can diminish the creative nature of coding projects, leading to only partial automation. Additionally, implementing nbgrader was laborious and was not completed due to deadlines. Future workshops or similar initiatives should try to implement tools, including AI-driven solutions, to automate and correct exercises, reducing the workload on the organizing committee.

#### 2. Enhancing educational strategies

Advancements in event structure and organization, along with improvements in didactic aspects, have been improved throughout editions. Daily feedback and final evaluations foster an environment open to change. Additionally, building confidence in new instructors with support from experienced teachers and incorporating accessibility concepts were also crucial and recommended approaches.

The use of Python as a high-level programming language, along with problem-solving exercises and cooperative learning techniques, is known to contribute to the improvement of students’ learning outcomes and motivation [44]. During the 2021 workshop, individual and group problem-solving exercises on biological issues led to complaints of overload (S3 Fig; S4 Table; S5 Table). In response, the 2022 edition focused solely on group exercises, reducing the study load and enhancing collaborative problem-solving and soft skills development.

An alternative strategy to improve programming skills, we used playful and didactic examples employing Graphics Interchange Format (GIFs) and quizzes (https://github.com/brazilpythonws/wpbd_update_2021_2022). This approach can help to make complex concepts more enjoyable and understandable for students.

Additionally, we incorporated computational thinking in lectures, focusing on abstraction and programming logic. Diversifying teaching strategies facilitates a better understanding of the content, given that student groups cover different educational backgrounds and demographic characteristics [45].

We prioritize developing Python skills in code construction, documentation, and modification, adhering to community standards [31]. To ensure increased usability, shareability, and reproducibility of materials, we strongly recommend the incorporation of Python’s best practices during live coding sessions, incorporating the guidelines provided in PEP8 [31].

The transition from a trainee to a trainer is facilitated by active participation in a network and the acquisition of proper training [24]. Our initiatives, including the Code Club, Train-the-Trainers-like event, and pair-teaching, created an engaging environment that boosts confidence in the coordination team to act as instructors or teaching assistants. Code clubs offered active and positive space for beginners and experienced members to enhance their Python and teaching skills [5]. The Train-the-Trainers-like event addressed instructors’ and assistants’ concerns, facilitating the transfer of experience and knowledge between former and current members.

While not explicitly required in any of the recent editions of the workshop, accessibility is crucial in developing introductory programming courses [46]. In recent editions of the event, we have made efforts to incorporate these topics to make the workshop materials more accessible and address various barriers, such as adapting figures of the ethics committee reports to accommodate color blindness. However, work remains to improve accessibility. Creating an inclusive environment requires a comprehensive understanding and application of accessibility, and this can be more easily achieved with the support of institutionalized assistance and the formation of specific committees for this purpose [47].

In this regard, accessibility concepts could be included from the selection and promotion of the event (enabling students with physical and/or learning disabilities to feel comfortable registering) to the training of the organizing committee (incorporating an understanding of concepts, and building knowledge and skills to ensure accessible classes and materials) [48,49]. In the immediate term, feasible perspectives for the event include introducing accessibility concepts in the Train-the-Trainers-like and Code Clubs and preparing teaching materials with these concepts in mind.

Another challenge faced by similar initiatives is democratizing and diversifying the audience and effectively including minoritized groups. In other words, what are the most suitable parameters used for this task, and what are the groups that should be considered? In Brazil, women’s underrepresentation in science is notable, with fewer women holding scholarships in well-funded areas and at higher levels [50]. Women’s presence also diminishes with career progression [51]. In this regard, our team opted to guarantee 50% of the spots to participants who declared themselves as a woman, as women are not a minority but rather a minoritized group [52], and to empower more women in programming careers (S4 Fig).

Our current strategy has some issues. For instance, it doesn’t adequately account for non-binary or other gender identities. Additionally, if the majority of enrollers are women, our approach could unintentionally maximize men’s participation to 50%. In future events or similar workshops, we recommend a system with exclusive vacancies for minority groups, avoiding these “side effects” and ensuring their correct inclusion. The remaining seats will be open to the general public, following the registration ranking, which can also include those groups.

For future workshops, we strongly recommend the inclusion of other minoritized groups among participants, speakers, and the organizing committee, such as underrepresented racial and ethnic groups. To achieve this, including a committee member with a degree in human sciences to guide decisions on these issues (S4 Fig) could be a valid strategy.

#### 3. Engagement and social media commitment

Following the COVID-19 pandemic, the demand for social media platforms for educational and scientific research activities has increased, especially for youth and adult education [35,53]. These platforms can showcase scientific discoveries, facilitate discussions, and promote critical thinking, thereby creating dynamic educational environments [54]. To enhance our impact, we have utilized popular social media platforms throughout the editions (Fig 3), increasing its reach and facilitating greater participant recruitment (S2 Fig). The accessible and collaborative space allowed us to share research and foster a community [55]. Despite these benefits, creating accessible, impactful content remains a challenge for researchers [55], especially on visually demanding platforms like Instagram. Training in design and engagement skills is needed, which can be supported through funding, as previously discussed. Therefore, we suggest supporting our community’s Python programming initiatives for bioscientists in Brazil through funding, experience exchange, and knowledge sharing.

### Updated recommendations for future workshop editions

The advances described in the previous section result from a lengthy process involving various editions and several organizing committees. Within this context, we present new recommendations (Table 2) that update those in Box 2 of [13] for students, researchers, and educators who wish to organize and promote events similar to ours.

**Table 2.** Updated recommendations for new workshop editions.

| **Structural topics** | **Focal point** | **Recommendations** |
| --- | --- | --- |
| Collaborative Forces in Structure | Workshop structure | Maintain an organized and clear core structure. It is important to delegate subjects to all groups and centralize problems with the Coordination team. |
|  | Code of Conduct | Establish a thorough Code of Conduct depicting guidelines for participants and organizers. |
|  | Coordination team training | Train the team since it has been helpful and it creates a sense of continuity/community. |
|  | Reliable feedback and improvements | Use virtual forms for reliable daily and overall event feedback. This step is essential to rethink strategies and implement new ideas. |
|  | Digital platforms | Use free, friendly, and collaborative digital platforms. Digital platforms are essential for coordination, didactics and training, and democratization, inclusion, and accessibility. |
|  | Programming best practices | Include specific slots for talking about coding best practices, but keep it simple if the event is designed for beginners. |
|  | Material sharing | Share the material used during the event with the community shortly after the end of the workshop, especially with those who attended it. Using a known share-coding platform (e.g., GitHub). Attributing a DOI (using e.g., Zenodo) to it is preferable. |
|  | Automated grading | Use AI-driven tools to automate and correct exercises, reducing the workload on the organizing committee. |
|  | Funding support | Consider securing funding support for at least one team member. The funded member can dedicate more time and resources to the event's planning, organization, and execution. We recommend organizing teams to apply for scholarships from national or institutional funding agencies. In addition, seeking sponsors is a good idea to provide rewards to the organizing team and the participants. |
|  | Research engagement (publication) | We recommend the coordination team to be involved in publishing, as it keeps them motivated and enhances their connection to the academic community. This engagement strengthens their connection with each other and stimulates creative thinking, which can enhance the overall quality of the event. |
| Enhancing Educational Strategies | Python training and support on teaching | Adopt a dual teaching approach, involving two instructors: one with more experience and the other with less. Additionally, establish a code club among the coordination members during the months preceding the event to study basic and intermediate Python coding. |
|  | Enhance accessibility | Adapt class material to increase the accessibility: use alt text, color-blind-friendly design, video conferencing platforms that support captions or offer transcripts, and translate disability or accessibility needs into student experience enhancements like sign language interpreters for in-person or hybrid events. |
|  | Select participants without personal bias | During the participant selection stage, apply filters that align with the expectations of the workshop scope, but do not provide the names and other information that could identify a candidate. |
|  | Prioritize diversity | Prioritize inclusivity by requesting gender, geographic location, etc., in the enrollment form, and leverage this information to enhance diversity during selection. |
| Engagement and Social commitment | Communication with the target audience | Establish a dedicated media and communication committee and maintain constant contact with the event's community. To facilitate interaction with the course's target audience and enhance the event's visibility, we recommend diversifying communication approaches in both scientific and non-scientific language. |
|  | Promotional Materials | Utilize free and collaborative online graphic design tools, as well as AI-powered tools, to create engaging, educational, and precise promotional materials for the event, thereby optimizing communication with the audience. |
|  | Increase visibility | Enhance the visibility of the event and its activities through regular dissemination on local and international scientific communication networks, particularly when research work is being developed. |
|  | Collaborative Network | Collaborate with other initiatives to establish a mutually beneficial network and amplify the impact and relevance of the event. Sharing information about similar groups and opportunities is a good starting point. |

## Conclusion

The Brazilian Python Workshop for Biological Data successfully incorporated suggestions and recommendations put forward by [13]. This manuscript highlights the progress made during the 2021 and 2022 editions and explores further suggestions for enhancing workshop experiences for bioscientists with limited or no programming background. Advancements were reported in the organizational structure of the event, democratization within the workshop, didactic approaches, programming best practices, scientific reproducibility, and social media and scientific dissemination. Despite these advancements, challenges have arisen that still require special attention in order to address them comprehensively. Additionally, we have provided an updated version of Box 2 from [13] for students, researchers, and educators interested in organizing and promoting events similar to ours.

## Ethics statement

The study was approved by the Ethics Committee on Animal Use (Protocol No. 06174/14) of FCAV/Unesp, Jaboticabal for the 2021 edition and was registered at the Brazilian Ethical Office (Plataforma Brasil: 45395321.0.0000.9029). The 2022 edition was approved by the Human Research Ethics Committee of ESALQ (Protocol No. 039771/2022) and was registered at the Brazilian Ethical Office (Plataforma Brasil: 58086522.2.0000.5395). All participants provided consent for both editions through an online form prior to the beginning of the workshop, and their confidentiality was ensured during data collection by replacing names with alphanumeric codes.

## Acknowledgements

Authors also thank research foundations that made this research possible and founded their research over a period overlapping the organization of at least one of the workshops: São Paulo Research Foundation (FAPESP), Coordination for the Improvement of Higher Education Personnel (CAPES), the Brazilian National Council for Scientific and Technological Development (CNPq) and Fundação de Estudos Agrários Luiz de Queiroz (FEALQ). RACdS, RB, MIGF and USB hold FAPESP scholarships (process numbers: 2021/11057-0, 2019/24412-2, 2023/16710-9 and 2023/09410-9, respectively). GSC and RMS hold CNPq scholarships (process numbers: 142289/2020-5 and 142262/2020-0, respectively). LGB and GSC hold a CAPES scholarship (process numbers 88887.802720/2023-00 and 88887.831628/2023-00, respectively). LGB and MIGF hold CAPES scholarship Finance code 001. GSC holds a FEALQ scholarship (project number 1059).

