## Supplementary Data for "Bridging the Python Training Gap for Bioscientists in Brazil: Improvements and Challenges"

**Institutions**

^1^University of São Paulo (USP), Campus Luiz de Queiroz, Department of Genetics, Genomics Group, Piracicaba, São Paulo, Brazil.

^2^Federal University of Ouro Preto (UFOP), Department of Biological Science, Ouro Preto, Minas Gerais, Brazil.

^3^São Paulo State University (UNESP), Vector-Borne Bioagents Laboratory, Department of Pathology, Reproduction and One Health, School of Agricultural and Veterinary Sciences, Jaboticabal, SP, Brazil.

^4^São Paulo State University/ School of Agricultural and Veterinary Sciences, Department of Exact Sciences, Jaboticabal, São Paulo, Brazil.

^5^University Of Guelph, Department of Animal Biosciences, N1G 2W1, Guelph, ON, Canada

^6^Free University of Berlin, Institute of Chemistry and Biochemistry, Berlin, Berlin, Germany.

^7^São Paulo State University (UNESP), Institute of Biosciences, Rio Claro, São Paulo, Brazil.

^8^Federal University of Pará (UFPA)/Francisco Mauro Salzano Molecular Biology Laboratory, Institute of Biological Sciences, Belém, Pará, Brazil.

^9^State University of Campinas/Department of Genetics, Evolution, Microbiology, and Immunology/Laboratory of Genomics and BioEnergy (LGE), Campinas, SP, Brazil.

^10^Laboratório de Biologia de Sistemas Regulatórios LABIS, Centro de Energia Nuclear na Agricultura, Universidade de São Paulo, Piracicaba, São Paulo, Brazil.

^11^University of São Paulo (USP) - Campus Luiz de Queiroz, Department of Genetics, Piracicaba, São Paulo, Brazil.

^12^São Paulo State University (UNESP), Department of Agricultural and Environmental Biotechnology, Varani's LAB, Jaboticabal, São Paulo, Brazil.

^13^Laboratory of Computational, Evolutionary, and Systems Biology, Centro de Energia Nuclear na Agricultura, Universidade de São Paulo, Piracicaba, São Paulo, Brazil.

^14^The Wallace Lab, Center for Applied Genetic Technologies (CAGT), University of Georgia, Athens, Georgia, United States of America.

**S1 Table. Brazilian introductory training initiatives focused on manipulating biological data using Python.**

| **Title** | **Institution** | **Is it paid?** | **Website** |
| --- | --- | --- | --- |
| Introdução à Bioinformática: aplicações em genômica e transcriptômica | Fio Cruz | No | [Introdução à Bioinformática: aplicações em genômica e transcriptômica - Latíssimo (fiocruz.br)](https://campusvirtual.fiocruz.br/gestordecursos/hotsite/cvf-node-30225-submission-2633/apresentaaao/2681) |
| BIOINFORMÁTICA – 60 HORAS \| CURSO CAPACITAÇÃO | IPB - Instituto Pedagogico Brasileiro | Yes | [Curso de Capacitação em Bioinformática - 60 horas (institutoipb.com.br)](https://www.institutoipb.com.br/cursos/cursos-de-capacitacao/bioinformatica/) |
| Introdução à Computação para Bioinformática | OnlineBioinfo/UFMG | Yes | [OnlineBioInfo (ufmg.br)](https://www.onlinebioinfo.dcc.ufmg.br/) |
| Algoritmos para Bioinformática | OnlineBioinfo/UFMG | Yes | [OnlineBioInfo (ufmg.br)](https://www.onlinebioinfo.dcc.ufmg.br/) |
| GBMeeting | Unicamp | No | [O EVENTO \| Gbmeeting (gbmeetingunicamp.wixsite.com)](https://gbmeetingunicamp.wixsite.com/gbmeeting) |
| Introdução à Linguagem Python aplicado a problemas Biológicos. | ITAP/EPAMIG | No | [Introdução Linguagem Python aplicado a problemas Biológicos. (epamig.br)](https://epamig.br/itap/cursos/introducao-a-bioinformatica-aplicada-a-agropecuaria/) |
| Curso de Verão em Bioinformática | USP | Yes | [Apolo (usp.br)](https://uspdigital.usp.br/apolo/apoObterCurso?cod_curso=460300035&cod_edicao=24001&numseqofeedi=1) |
| Curso de Férias em Bioinformática | UFMG | Yes | [Home (bioinfo.com.br)](https://www.ufmg.bioinfo.com.br/) |
| Escola Latino-Americana de Bioinformática | LNCC | No | [Escola Latino-Americana de Bioinformática - ELAB/LNCC](https://www.elab.lncc.br/) |
| Escola Gaúcha de Bioinformática | UFRGS | Yes | [Home - EGB - Escola Gaúcha de Bioinformática (ufrgs.br)](https://www.ufrgs.br/egb/) |
| Escola Paranaense de Bioinformática | UEL/UFPR/UTFPR | Yes | <https://www.even3.com.br/epbioinfo2024/> |
| Workshop de Python para dados de Biologia Vegetal |  | No | <https://python4plantdatabr.wixsite.com/wspythonplantbio2022> |
| Workshop de Python para dados de Microbiologia | UFCSPA | No | <https://python4microbiobrazil.gitlab.io/sitepythonmicrobiodata2022/> |
| From Gene to Trait | GCCRC (Unicamp/Embrapa) | No | https://www.gccrc.unicamp.br/course-from-gene-to-trait/ |

**S2 Table. Schedule of the Brazilian Python Workshop for Biological Data in 2021.**

| **Time** | **Day 1** | **Day 2** | **Day 3** | **Day 4** | **Day 5** |
| --- | --- | --- | --- | --- | --- |
| 8h30 | Welcome | Presentation: Introduction to Pandas and Matplotlib | Reserved for group and individual Exercises, questions and answers | Presentation: Results obtained in the group exercises | Presentation: Problem of the Day |
| 9h00 | Live Coding - Introduction to Python and its data structures | Live Coding – Introduction to Tabulated Data Manipulation |  |  | Live Coding - Exploration of biological diversity data |
| 10h00 |  |  |  | Live Coding – Exploration of epidemiological data |  |
| 10h30 |  |  |  |  |  |
| 11h00 |  |  |  |  |  |
| 12h00 | Break | |  | Break | |
| 13h00 | Talk: MitoHifi – finding and annotating circular molecules | Talk: Machine learning applied to biological issues |  | Talk: Analysis of biological networks in Python | Talk: Best practices and reproducibility of Python scripts |
| 14h00 | Live Coding - Introduction to Python and its data structures | Live Coding – Graphical Exploratory Data Analysis |  | Live Coding – Exploration of epidemiological data | *Flash Talks* - Participants Projects |
| 15h00 |  |  |  |  | Live Coding - Exploration of biological diversity data |
| 16h00 |  |  |  |  |  |
| 17h00 - 18h00 | Questions and Answers | |  | Questions and Answers | Workshop Closure |

**S3 Table. Schedule of the Brazilian Python Workshop for Biological Data in 2022.**

| **Time** | **Day 1** | **Day 2** |  | **Day 3** | **Day 4** |
| --- | --- | --- | --- | --- | --- |
| 8h30 | Welcome | Presentation: Introduction to Python and its data structures | Reserved for group Exercises, questions and answers | Presentation: Results obtained in the group exercises | Presentation: What is Biopython? |
| 9h00 | Live Coding - Introduction to Python and its data structures |  |  |  | Live Coding - Introduction to Biopython |
| 10h00 |  | Coffee-break |  | Live Coding - Autonomy for biological data analysis in Python |  |
|  |  | Live Coding - Introduction to Pandas and Matplotlib |  |  |  |
| 11h00 |  |  |  |  |  |
| 12h00 | Break | |  | Break | |
| 13h00 | Talk:  Eco-Evolutionary Simulation in Python | Talk: Running external programs with Subprocess lib |  | Talk: Best practices and reproducibility of Python scripts | Talk: What is possible to do with python basics? |
| 14h00 | Live Coding - Introduction to Python and its data structures | Live Coding - Introduction to Pandas and Matplotlib |  | Live Coding - Autonomy for biological data analysis in Python | Flash Talks - Participants Projects |
| 15h00 |  |  |  |  | Live Coding - Introduction to Biopython |
|  |  | Coffee-break |  |  |  |
| 16h00 |  | Live Coding - Introduction to Pandas and Matplotlib |  |  |  |
| 17h00 - 18h00 | Questions and Answers | |  | Questions and Answers | Workshop Closure |

**S1 Text. Code-club topics explored by the 2022 organizing team.**

1. PEP8; Basic Syntax and Variables.
2. Numeric operations (integers, floats, complex numbers); Comparison, Logical, and Boolean Operators
3. Indexing; methods and functions applied to Strings
4. Sequences (Tuples, Lists, Ranges)
5. Conditional statements (if, else, elif)
6. Sets
7. Loops (for, while)
8. Dictionaries
9. Functions and Recursive Functions
10. Input and Output formatting, and Import
11. NumPy
12. Data Manipulation with Pandas - Series
13. Data Manipulation with Pandas - DataFrame
14. Data Manipulation with Pandas - File Import and Exploration with Different Methods; exploratory statistical analyses
15. Data Cleaning and Preparation
16. Data Visualization with Matplotlib (parte 1)
17. Data Visualization with Matplotlib (parte 2)

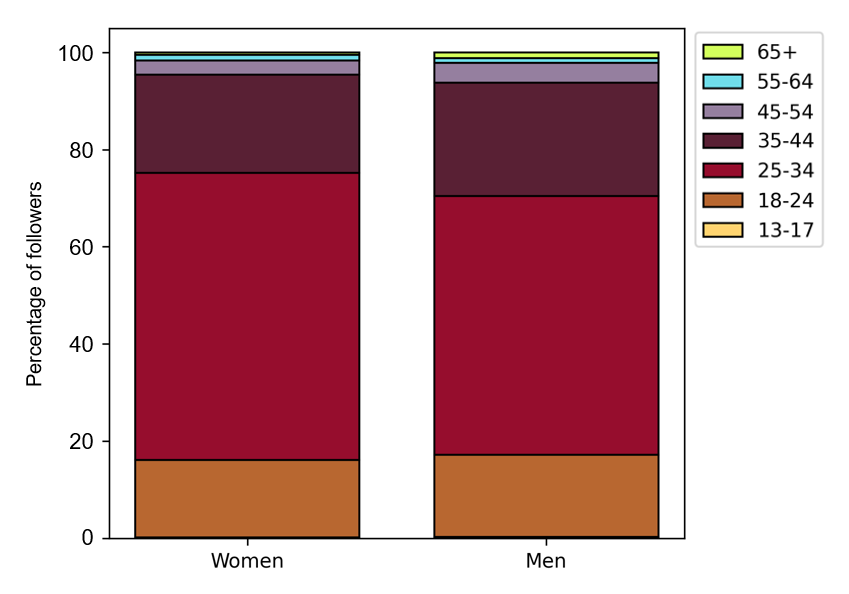

**S1 Fig. Age characterization of Instagram followers of the Brazilian Python Workshop for Biological Data.** The age distribution between male and female followers of the Instagram page for the Brazilian Python for Biological Data Workshop (in 14th September, 2023).

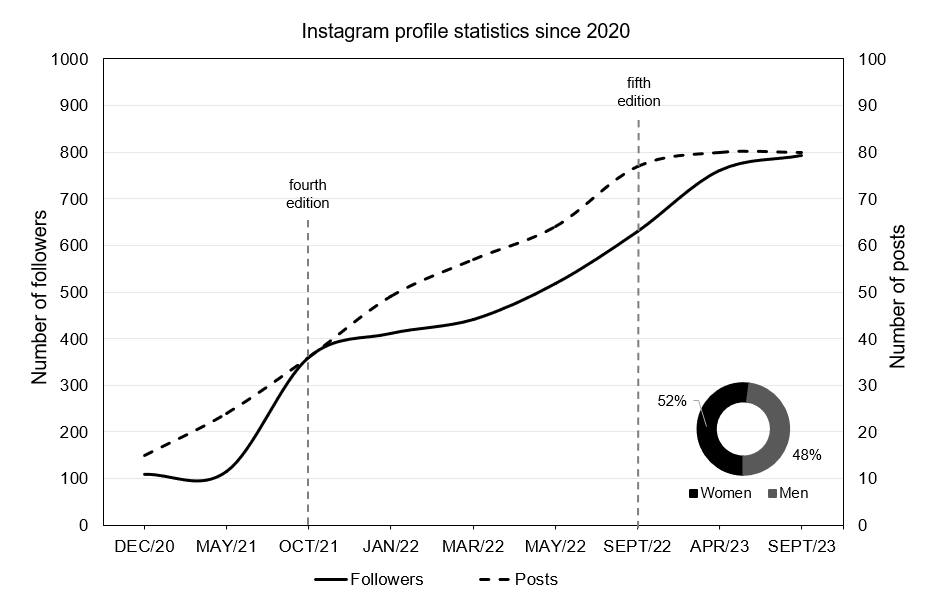
**S2 Fig. The Brazilian Python Workshop for Biological Data Instagram profile statistics since third edition (2020).** The evolution of the number of followers on the Instagram profile of our workshop, along with the evolution of the number of posts made by the same profile. The donut plot shows the distribution between male and female followers of the workshop's Instagram page (until 14th September, 2023).

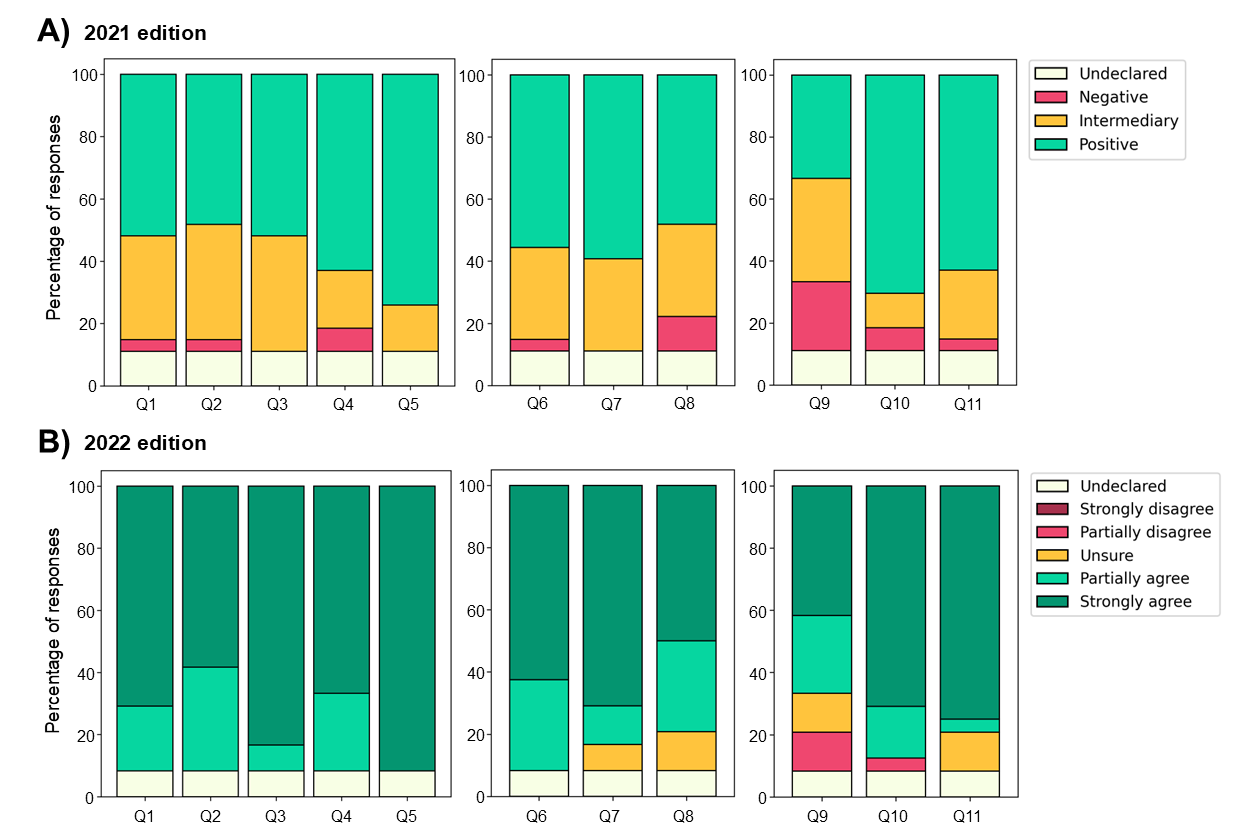
**S3 Fig. Course structure evaluation about** (A) Evaluation of instructors and presentations, where Q1 = The instructors were effective, Q2 = The presentations were clear and organized, Q3 = The instructors fostered student engagement, Q4 = The instructors managed their time effectively during the classes, and Q5 = The instructors were accessible and helpful.; (B) The objectives and materials presented, where Q1 = The objectives were clear, Q2 = The course content was organized and well-planned, and Q3 = The instructional material was well-developed.; (C) The organization and the tools used, where Q4 = The course workload was appropriate, Q5 = The tools were suitable for the course, and Q6 = The course was structured to enable the participation of all students.

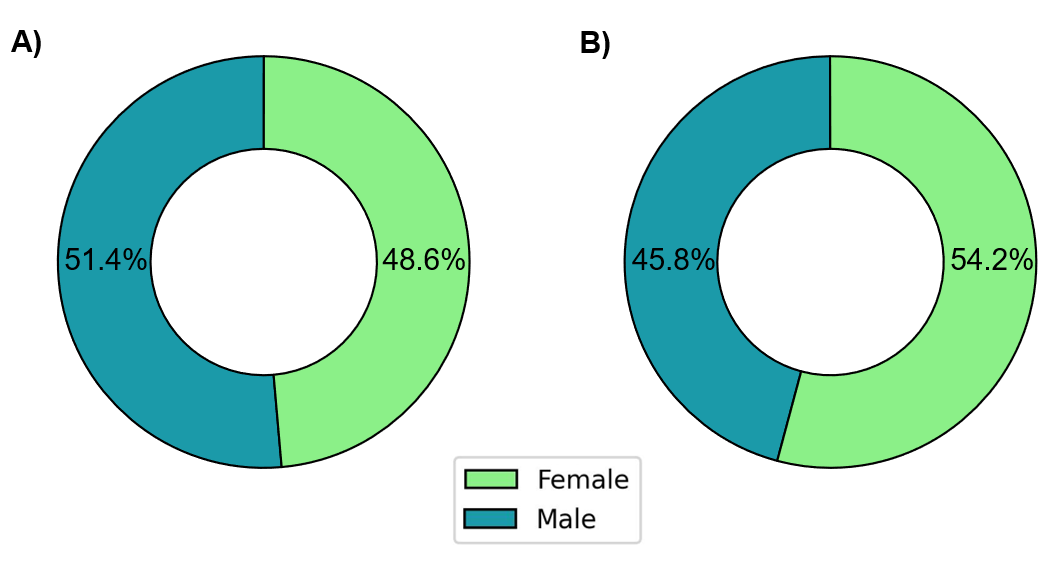

**S4 Fig. Gender ratio among selected participants.** Distribution of gender among participants in the 2021 edition (A) and the 2022 edition (B).

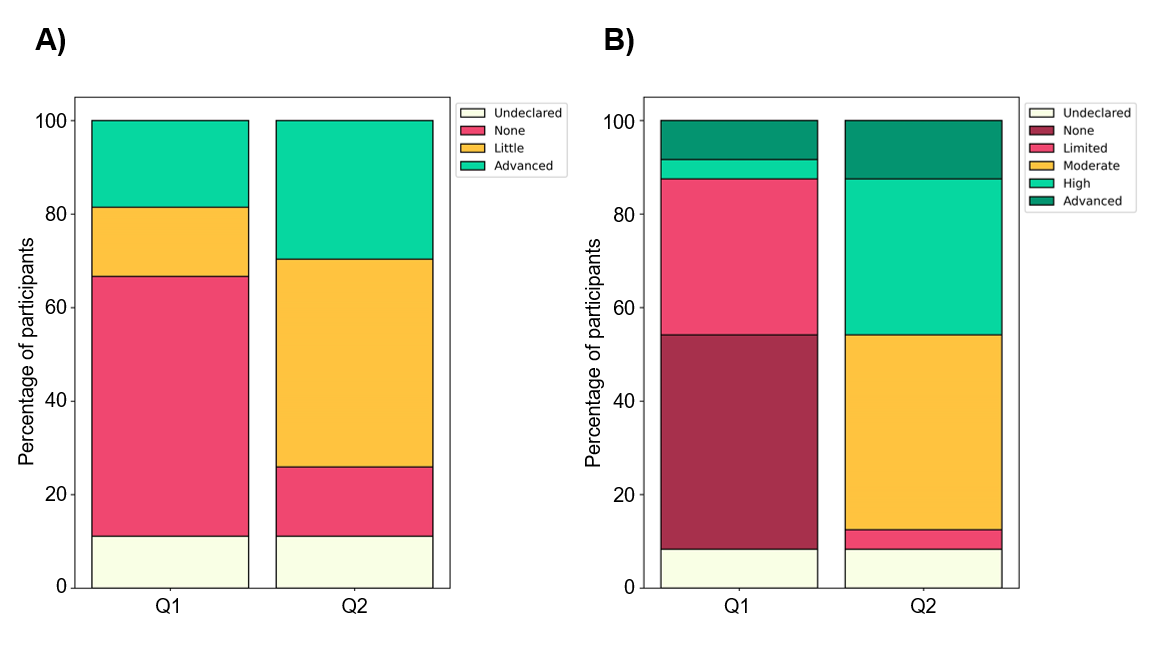

**S5 Fig. Participants' Knowledge of the Python Programming Language.** Changes in participants' knowledge of the Python programming language, where Q1 = Programming knowledge level at the beginning of the course and Q2 = Programming knowledge level at the end of the course.

**S4 Table. Representative answers obtained from participants at the end of the IV Python for Biological Data Workshop, held in 2021.**

| **Question** | **Category** | **Answer** |
| --- | --- | --- |
| How would you improve this course? | Making the notebooks available | "I think making the notebooks available at the end of the course could be interesting for those who are learning and missed some parts." |
|  |  | "Providing the notebooks with the practical activity codes ready, and commenting on the codes during the class. This would help students focus on the explanation and prevent constant interruptions due to difficulties in typing the codes and following the explanation simultaneously." |
|  |  | "I would like to suggest providing an outline of the topics that will be covered each day. If the class recording and the notebook are not made available, perhaps a practice outline for the student to follow later could help." |
|  | Workload | "I believe the schedule could be better distributed. I found it heavy to have just one day dedicated to group and individual activities." |
|  |  | "Better organizing the day for individual and group exercises. Having both due on the same day was detrimental to the performance of at least one of them." |
|  |  | "Having more time for the exercises (extended deadlines) felt tight." |
| What was your experience participating in the distance learning course? Would you take another course in the same format? | Positive experience | "I enjoyed the experience in the distance learning course. I don't see myself being able to travel to the event location if it were in another format. I would definitely attend another event in this format." |
|  |  | "I found the experience very good. The organizers managed to adapt the course very well to the virtual environment." |
|  |  | "It was very good. I would definitely do another one." |
|  | Neutral | "It was interesting, I think it could be spread over more days because the hours felt a bit heavy." |
|  |  | "Although I prefer the in-person format, I believe that adopting Slack + Meet + Google Colab adequately covered the course's needs, and I think the remote format is efficient for its execution." |
|  |  | "It was normal, I'm already accustomed to the pace of distance learning courses. Yes." |
|  | Negative experience | "[...] I believe that the learning in this course was hindered by the online method [...], as it was very difficult to follow the teacher's explanation and at the same time copy the code [...] Maybe I would take another online course, but only if I had a second screen available to connect to my notebook. However, I will definitely prefer in-person courses." |

The responses contained in the feedback forms were translated from the original language (Portuguese).

**S5 Table. Representative answers obtained from participants at the end of the 5th Python Workshop for Biological Data, held in 2022.**

| **Question** | **Category** | **Answer** |
| --- | --- | --- |
| How would you improve this course? | Making the notebooks available | "The scripts should be made available because it's difficult to listen to the explanation, type the code, run it, encounter errors, and then go to Discord. It's very easy to get lost and fall behind." |
|  |  | "Providing the Colabs and study material in advance for study before the lecture." |
|  |  | "Provision of handouts on the content of the classes, for prior study. I think it would be better for keeping up." |
|  | Workload | "Very extensive workload, sometimes it becomes tiring due to other tasks in our daily lives, and more time for lunch." |
|  |  | "More days dedicated to Biopython (minimum 2 days)." |
|  |  | "[...] perhaps distribute the workload better." |
| **Which aspects of this course were most useful or valuable to you?** | Content | "Learning to use new tools for data manipulation." |
|  |  | "The part about creating graphs, like box plots." |
|  |  | "Understanding the programming logic related to each method/function." |
|  | Methodology | "The entire structure of the course, from the lectures to the workshops, exercise resolutions, and flash talks. It was a very valuable first contact." |
|  |  | "How to use Python, searching for what you don't know, and the introduction to the main libraries." |
|  |  | "The applicability of Python in projects developed in the field of biology." |
|  | Networking | "[...] In terms of value, real-time learning, networking, working with peers to develop activities." |

The responses contained in the feedback forms were translated from the original language (Portuguese).
